# No more business as usual: agile and effective responses to emerging pathogen threats require open data and open analytics

**DOI:** 10.1101/2020.02.21.959973

**Authors:** Galaxy and HyPhy developments teams, Anton Nekrutenko, Sergei L Kosakovsky Pond

## Abstract

The current state of much of the Wuhan pneumonia virus (COVID-19) research shows a regrettable lack of data sharing and considerable analytical obfuscation. This impedes global research cooperation, which is essential for tackling public health emergencies, and requires unimpeded access to data, analysis tools, and computational infrastructure. Here we show that community efforts in developing open analytical software tools over the past ten years, combined with national investments into scientific computational infrastructure, can overcome these deficiencies and provide an accessible platform for tackling global health emergencies in an open and transparent manner. Specifically, we use all COVID-19 genomic data available in the public domain so far to (1) underscore the importance of access to raw data and to (2) demonstrate that existing community efforts in curation and deployment of biomedical software can reliably support rapid, reproducible research during global health crises. All our analyses are fully documented at https://github.com/galaxyproject/SARS-CoV-2.

---

The initial publications describing genomic features of COVID-19 [1–4] used Illumina and Oxford nanopore data to elucidate the sequence composition of patient specimens (although only Wu et al. [3] explicitly provided the accession numbers for their raw short read sequencing data). However, their approaches to processing, assembly, and analysis of raw data differed widely (Table 1) and ranged from transparent [3] to entirely opaque [4]. Such lack of analytical transparency sets a dangerous precedent. Infectious disease outbreaks often occur in locations where infrastructure necessary for data analysis may be inaccessible or unbiased interpretation of results may be politically untenable. As a consequence, there is a global need to ensure access to free, open, and robust analytical approaches that can be used by anyone in the world to analyze, interpret, and share data. Can existing tools and computational resources support such a global need? Here we show that they can: we analyzed all available raw COVID-19 data to demonstrate that analyses described in [1–4] can be reproduced on public infrastructure using open source tools by any researcher with an Internet connection.

**Table 1.** Methods used for the analysis of primary COVID-19 data. ? = uncertain (e.g., Holshue et al. [4] identify FreeBayes as an assembly tool).

| Analysis stage | Publication |  |  |  |
| --- | --- | --- | --- | --- |
|  | [3] | [1] | [2] | [4] |
| Tools | Bowtie2<br>MegaHit<br>Trinity<br>MAFFT<br>PhyML<br>MEGA<br>RDP4 | BWA<br>Geneous<br>MegaHit<br>MAFFT<br>Clustal<br>RAxML | BWA<br>SPAdes<br>CNCBio<br>MAFFT<br>RAxML | minimap<br>Sequencher<br>FreeBayes? |
| Versions specified | + | + | + | + |
| Parameters specified | - | - | - | - |
| Raw data | + | ? | - | - |

We exclusively used free software tools publicly available from the BioConda package distribution system [5], deployed through the worldwide network of open Galaxy platforms [6] and executed using public high throughput computational infrastructure (XSEDE in the US, VSC in Belgium, de.NBI and ELIXIR in EU, NeCTAR Research Cloud in Australia). We also used an open source Jupyter environment [7] for exploratory analysis of data. All analyses performed here are fully documented and accessible at https://github.com/galaxyproject/SARS-CoV-2/ and https://doi.org/10.5281/zenodo.3685264 (note that these are being continuously updated).

We divided our analysis into the following stages: (1) read pre-processing, (2) genome assembly, (3) timing the most recent common ancestor (MRCA), (4) analysis of genomic variation within individual samples, and (5) recombination and selection analyses.

We pre-processed six currently available (as of Feb 19, 2020) sequencing read datasets for COVID-19 (Table S1) by removing adapter contamination and reads derived from human transcripts and combined the resulting datasets. This was done to COVID-19-specific reads that constitute only a fraction of the original data. These were used as inputs for SPAdes assembler [8] and Unicycler [9]—an assembly pipeline based on SPAdes that includes a number of pre-processing and polishing steps. Both approaches were able to reconstruct a full length COVID-19 genome with Unicycler producing a cleaner assembly graph. Its largest contig (29,781bp) had 100% identity to the published assembly NC_045512.

Next we estimated the date of the most recent common ancestor (MRCA) of COVID-19. For this we used simple root-to-tip regression [10] (more complex and powerful phylodynamics methods could certainly be used, but for this data with very low levels of sequence divergence, simpler and faster methods suffice). Using a set of sequences from all COVID-19 sequences available as of Feb 16, 2020 we obtained an MRCA date of Oct 24, 2019, which is close to other existing estimates [11].

The vast majority of COVID-19 genomic data available at the time of writing are partially or fully assembled genomes. There is no public access to sequence reads that were used to produce these assemblies: as of Feb 19, 2020—more than two months since the beginning of the outbreak—there are only six raw datasets (Table S1).

This should be unacceptable, since raw read data can be used to uncover viral diversity within individual samples and to evaluate robustness and reliability of the assembly. To demonstrate that such diversity exists, we mapped Illumina reads against COVID-19 reference (NC_045512) and identified sequence variants with frequencies above 5% while taking into account quality of alternative bases and strand bias. Five percent was selected as a conservative threshold that can be reliably resolved from Illumina data [12]. Using this threshold, thirty nine single nucleotide variants (SNVs) were identified in total across all samples (Fig. 1). The most prominent sequence variant was observed in sample SRR10903401. It is an A-to-C substitution with alternate allele frequency of 38% that causes a Lys^921^Gln amino acid replacement within the spike glycoprotein S (product of gene *S*). S is a homotrimeric protein containing S1 and S2 subdomains mediating receptor recognition and membrane fusion, respectively [13]. S2 subdomains contain two heptad repeat (repeats of units containing seven amino acids) regions: HR1 and HR2. The Lys^921^Gln substitution we observed is located in HR1 and forms a salt bridge with Gln^1188^ within HR2. This is one in a series of salt bridges involved in the formation of the HR1/HR2 hairpin structures [14]. This site invariably contains Lys in all human SARS-related coronaviruses (S protein residue 903) as well as in many other coronaviruses (Fig. 2). However, more distantly related coronaviruses including transmissible gastroenteritis coronavirus (TGEV), the porcine respiratory coronavirus (PRCV), the canine coronavirus (CCV), the feline peritonitis virus (FIPV), and the porcine epidemic diarrhea virus (PDEV) all contain Gln at the corresponding position ([14] and Fig. 2). The Lys^921^Gln change would prevent the formation of the salt bridge with Gln^1188^ and may have structural and functional implications for the spike protein structure and, consequently, COVID-19 virulence. This potentially adaptive change was not observed in the other two samples and lack of raw read data prevented us from identifying it in other geographically and temporally distributed samples.

**Figure 1.**
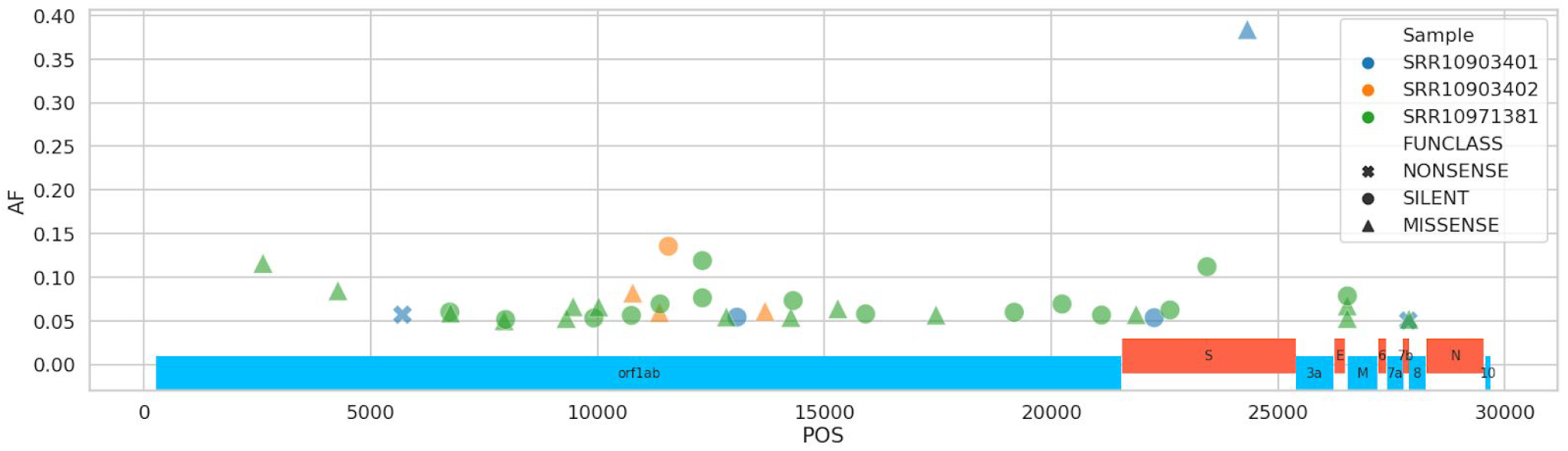
Distribution of nucleotide changes across COVID-19 genome. AF = minor allele frequency, POS = position

**Figure 2.**
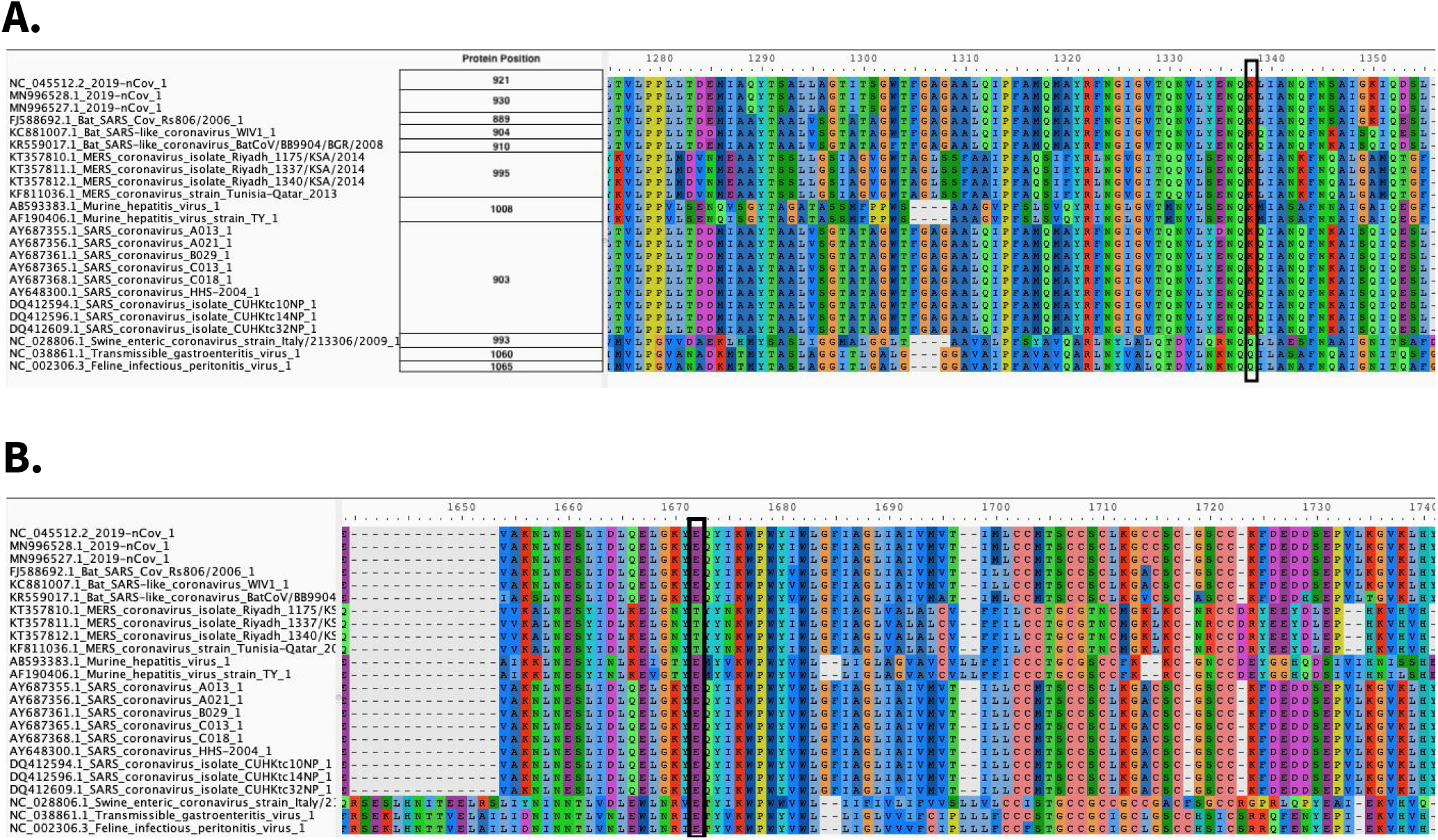
Amino acid alignment of spike glycoprotein regions HR1 (A) and HR2 (B). The site of the Lys^921^Gln substitution observed by us in a COVID-19 isolate is highlighted with a black rectangle in panel A. Its corresponding salt bridge partner is highlighted with black rectangle in panel B.

To detect potential genome rearrangement events that might have led to the emergence of COVID-19 we performed analysis of recombination using a genetic algorithm approach [15]. Wu et al. [3] identified two potential recombination breakpoints within the COVID-19 S-gene with some segments having higher similarity to Bat ZC45 and ZXC21 coronaviruses (accessions MG772933 and MG772934, respectively), while others were more similar to SARS Tor2 and SZ3 isolates (accessions AY274119 and AY304486). Our attempt at reproducing this analysis did identify a set of potential breakpoints similar to the ones reported by Wu et al. [3], but lacking robust statistical support (Fig. 3).

**Figure 3.**
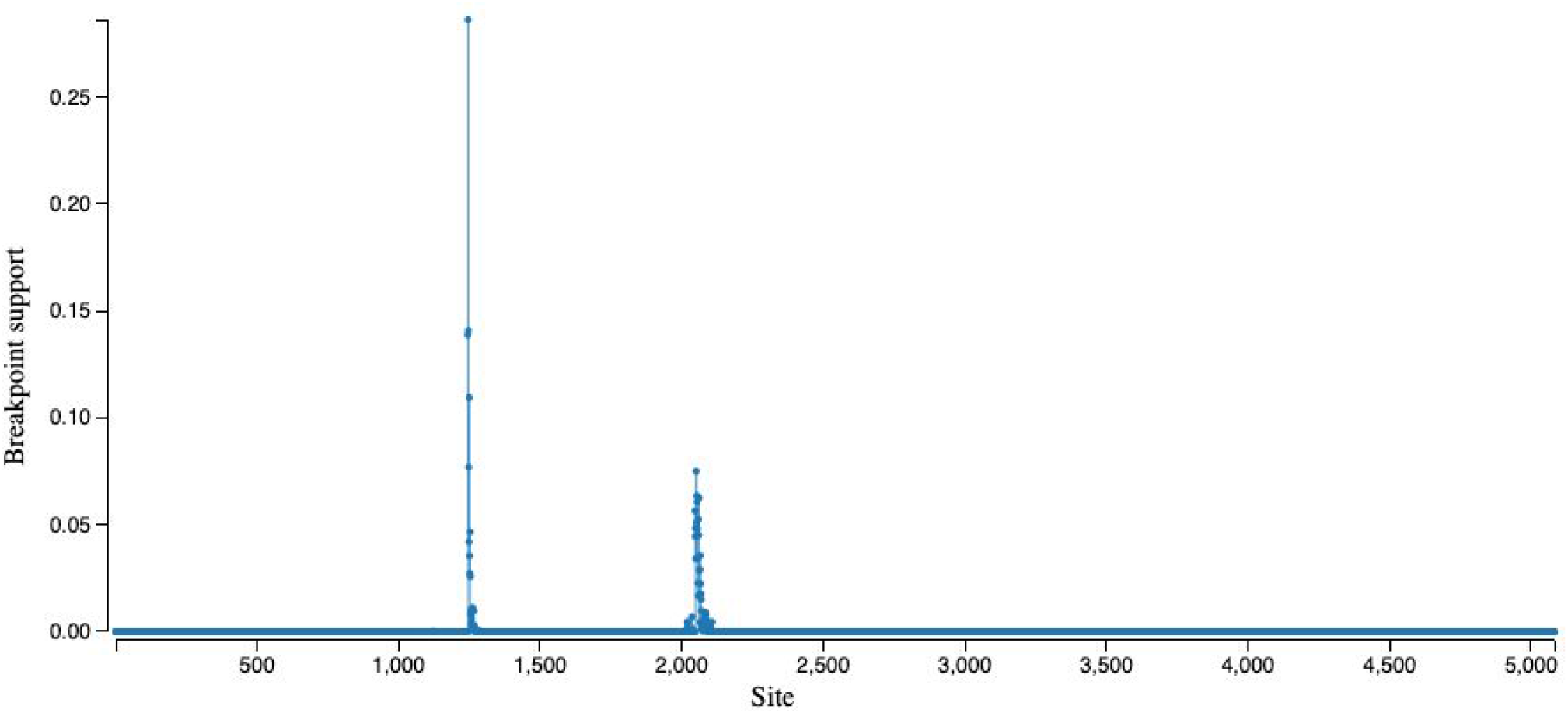
Location of potential recombination breakpoints along the *S-*gene (GARD analysis).

Finally, we performed a branch-level test for positive selection on a codon-alignment of the *S* gene from COVID-19, SARS-Tor2 as well as Bat ZC45, ZXC21, and Rp3 coronaviruses, specifically to identify if there was any evidence of diversifying selection along the ancestral branch leading to COVID-19 isolates. We found statistically significant evidence of positive diversifying selection (~7% of *S*-gene sites) along the branch leading to COVID-19 (Fig. 4).

**Figure 4.**
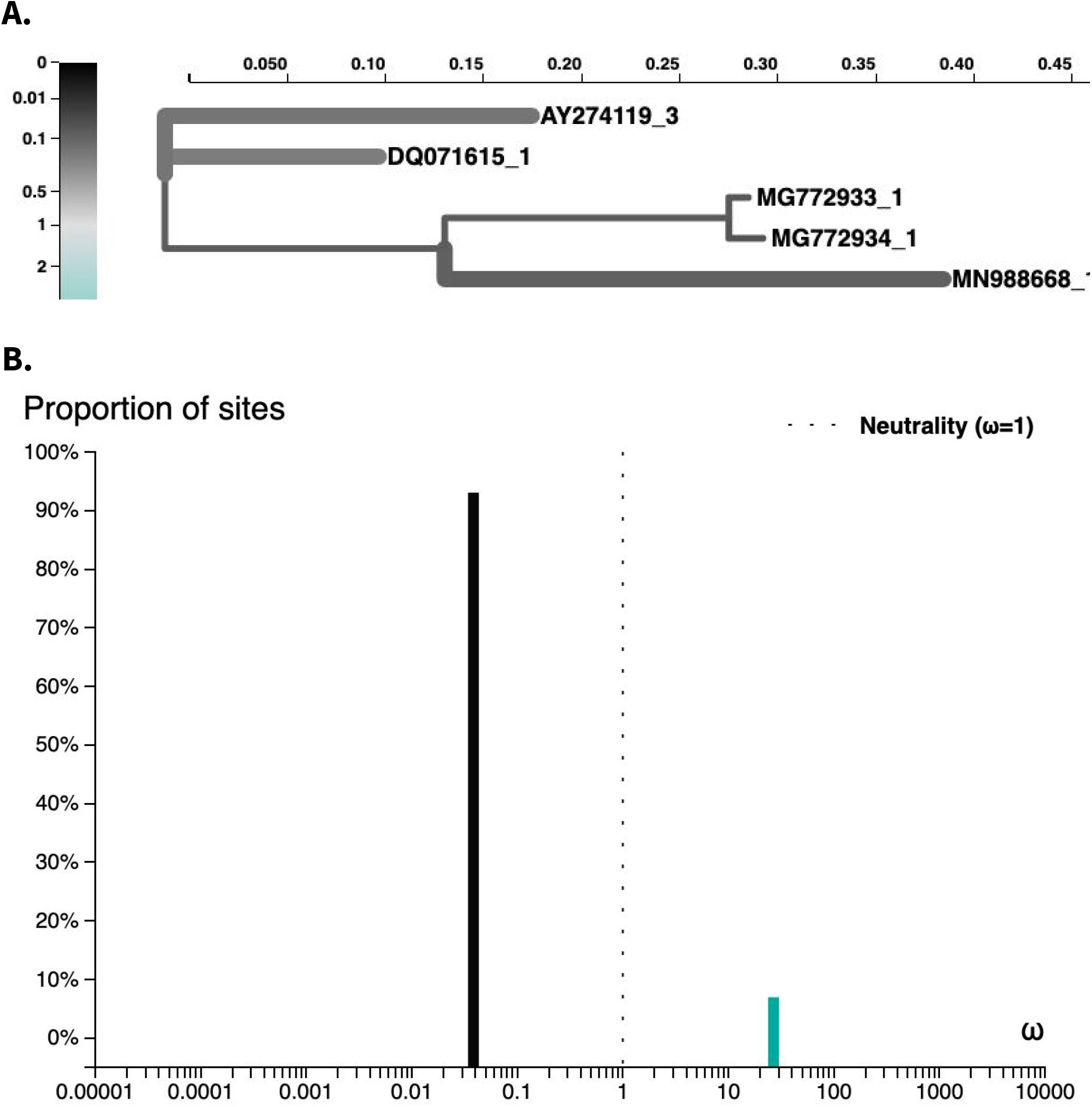
Analysis of branch-specific positive diversifying selection (aBSREL) along the branch leading to COVID-19 (MN988688).

The goal of our study was to (1) raise awareness of the lack of primary data necessary to effectively respond to global emergencies such as the COVID-19 outbreak and (2) demonstrate that all analyses can be performed transparently with already existing open source publicly available tools and computational infrastructure. The first problem—reluctance to share primary data—has its roots in the fact that the ultimate reward for research efforts is a highly cited publication. As a result, individual researchers are naturally averse to sharing primary data prior to manuscript acceptance. The second issue—underutilization of existing, community supported tools and analysis frameworks—may be due to the lack of sustained efforts to educate the biomedical community about best practices in (genomic) data analysis. Such efforts exist (e.g., [16]) but have difficulties reaching a wide audience because prominent scientific publication outlets are reluctant to accept data analysis tutorials or reviews. Yet the only way to improve accessibility and reproducibility of biomedical research is through dissemination of best analysis practices.

We want to particularly emphasize the issue of irreproducibility. All researchers involved in any given outbreak research should have access to a set of community-curated tools in the same way as they have access to COVID-19 RT-PCR primers [17]. Moreover they should have access to computational infrastructure that can execute these tools and apply them to potentially large NGS datasets. This is essential as precious time is spent on “reinventing the wheel”. Instead, in an ideal world, after reading any of the original COVID-19 manuscripts any researcher should be able to apply the same analytical procedures to their own data. To illustrate these points we assessed the reproducibility of the four initial manuscripts describing the COVID-19 genome (Table 2). All manuscripts reported versions of the software used, but none listed parameters used. This effectively prevents quality control and replication because outcomes of complex procedures such as genome assembly, phylogenetic reconstruction, and recombination analysis are notoriously parameter-dependent. One of the manuscripts [4] explicitly lists FreeBayes [18], a variant discovery tool, as software used for short read assembly—something that FreeBayes is not capable of doing. Finally, only [3] provided access to the raw data, rendering the other three manuscripts unverifiable and irreproducible.

Our short study demonstrates that viral genome analyses can be performed using open worldwide scientific infrastructure, relying entirely on community-curated and -supported open-source software. While we used Galaxy as the platform to execute all analyses described here, the individual software components can be obtained directly from BioConda and run independently. They can be combined into workflows using systems like the CWL[19], Nextflow[20], or Snakemake[21]. Whatever the execution environment or workflow engine, using community supported, versioned, open source tools makes data analyses robust and transparent. This increases the quality, efficiency and, ultimately, impact of biomedical research.

As we were submitting this manuscript, several new sets of raw Illumina reads have been deposited to the Short Read Archive at the NCBI. We did not analyze them in this version of our (Feb 23) manuscript. However, this reinforces the key point we are making in this study—anyone can use the open workflows described here to analyze the new data. In an age of digital connectedness, open, highly accessible, globally shared data and analysis platforms have the potential to transform the way biomedical research is done, opening the way to ‘global research markets’, where competition arises from deriving understanding rather than access to samples and data. Other disciplines have embraced the benefits of global data generation and sharing, astronomy and high energy physics being two highly successful examples. We have the opportunity to mirror their successes in infrastructure funding by demonstrating that biological research can embrace the same global perspective on common infrastructure investment and data sharing.

## Authors and contributions

Dannon Baker^1^, Marius Van Den Beek^2^, Daniel Blankenberg^3^, Dave Bouvier^4^, John Chilton^5^, Nate Coraor^6^, Frederik Coppens^7^, Ignacio Eguinoa^8^, Simon Gladman^9^, Björn Grüning^10^, Delphine Larivière^11^, Andrew Lonie^12^, Sergei Kosakovsky Pond^13^, Wolfgang Maier^14^, Anton Nekrutenko^15^, James Taylor^16^, Steven Weaver^17^

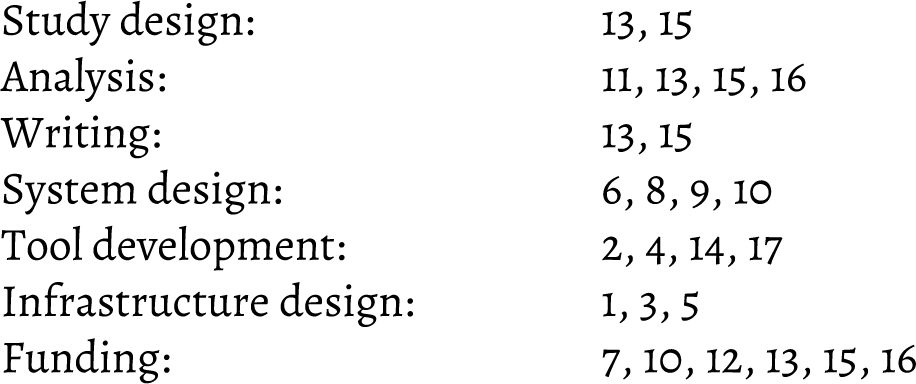

## Acknowledgments

Usegalaxy.org efforts are funded by NIH Grants U41 HG006620 and NSF ABI Grant 1661497. Usegalaxy.eu is supported by the German Federal Ministry of Education and Research grants 031L0101C and de.NBI-epi. Galaxy and HyPhy integration is supported by NIH grant R01 AI134384. Usegalaxy.org.au is supported by Bioplatforms Australia and the Australian Research Data Commons through funding from the Australian Government National Collaborative Research Infrastructure Strategy. Hyphy.org development team is supported by NIH grant R01GM093939. Usegalaxy.be is supported by the Research Foundation-Flanders (FWO) grant I002919N and the Flemish Supercomputer Center (VSC).

---

**Table S1.**
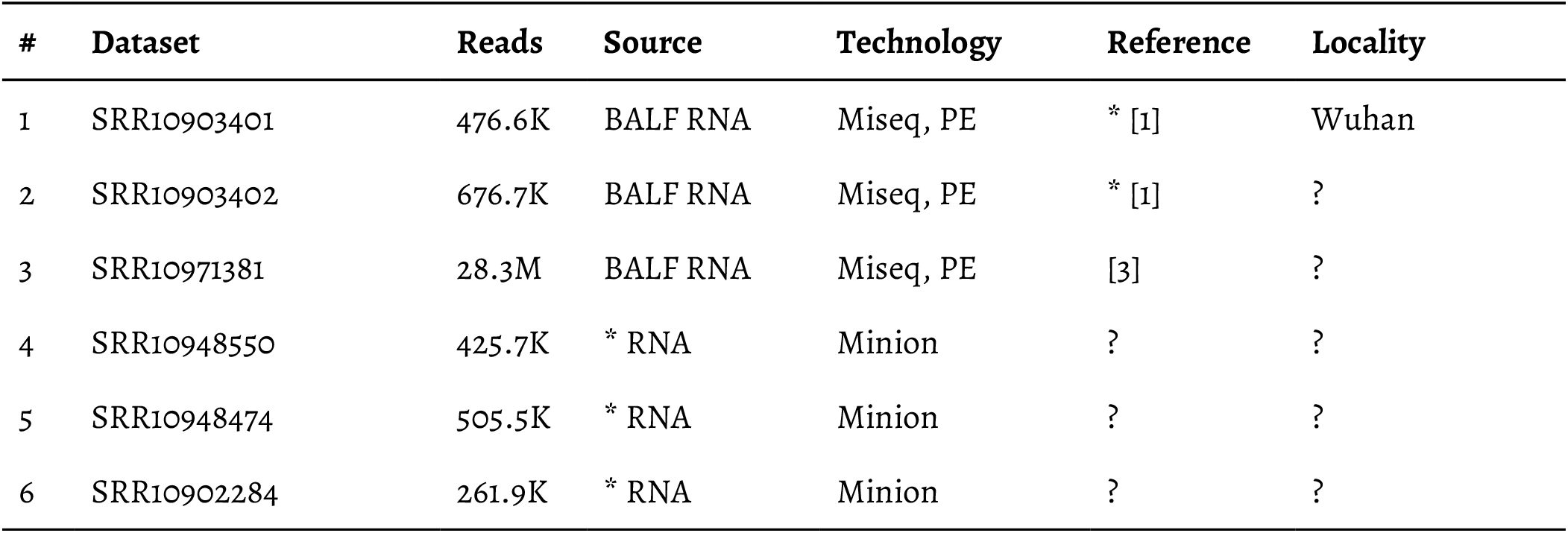
Raw COVID-19 sequencing data available at the time of writing (Feb 20, 2020). BALF = bronchoalveolar lavage fluid, * indicates that data may not be reliable (for example, the link between SRR10903402 and [1] is inferred: neither the SRA record nor the manuscript establish this relationship).

| # | Dataset | Reads | Source | Technology | Reference | Locality |
| --- | --- | --- | --- | --- | --- | --- |
| 1 | SRR10903401 | 476.6K | BALF RNA | Miseq, PE | * [1] | Wuhan |
| 2 | SRR10903402 | 676.7K | BALF RNA | Miseq, PE | * [1] | ? |
| 3 | SRR10971381 | 28.3M | BALF RNA | Miseq, PE | [3] | ? |
| 4 | SRR10948550 | 425.7K | * RNA | Minion | ? | ? |
| 5 | SRR10948474 | 505.5K | * RNA | Minion | ? | ? |
| 6 | SRR10902284 | 261.9K | * RNA | Minion | ? | ? |

